# Microfluidic Immunocapture Device for Direct Detection of Lyme Disease

**DOI:** 10.64898/2025.12.14.694202

**Authors:** Kyle Wellmerling, Brian Kirby

## Abstract

Lyme Disease is a multisystem infectious disease caused by the *Borellia burgdorferi* complex, and is a growing threat to public health. Approximately 476,000 people are infected with Lyme in the United States each year. Although Lyme is readily treated with antibiotics when detected early, early detection remains difficult. Current testing remains difficult because the standard 2-tiered ELISA/Western assay indirectly detects Lyme via measurement of a host immune response, which suffers from an inherent time-lag in host antibody production. A direct test for Lyme Disease would overcome these inherent limitations. To this end we report on the first microfluidic immunocapture device for Lyme Disease. We engineered a geometrically enhanced differential immunocapture (GEDI) technology to capture whole-organism *Borrelia* for direct on-chip detection. This approach is potentially amenable with other work in the field to develop direct PCR or aptamer tests for Lyme Disease, as our device could serve as a platform to drastically enhance the concentration of present *Borrelia* into a small volume.

## 1 Introduction

Lyme disease is a vector-borne disease caused by the bacterial spirochete *Borellia burgdorferi*. Lyme is transmitted to humans by Ixodid ticks, primarily by the Ixodes scapularis deer tick^1^. Based on experimental studies in animals^2^ and humans^3^, an infected tick must feed for 48-72 hours in order to transmit *Borellia burgdorferi* to the host human. After initial infection, Lyme disease manifests in three stages: early localized, early disseminated, and late disease^1^. The early localized disease stage is characterized by the appearance of the erythema migrans rash (single or multiple) at the site of the tick bite 3 to 30 days (typically 7 to 14) after the bite. The erythema migrans (single or multiple) is found in about 90% of patients with objective evidence of infection with *Borellia burgdorferi*^1,4,5^, and may be accompanied by systemic findings such as fever, malaise, headache, regional lymphoadenopathy, stiff neck, myalgia, and/or arthralgia. Early disseminated Lyme disease occurs after dissemination of *Borrelia* to distal sites, both through hematogenous^6^ and non-hematogenous (lymphatic system or direct spread through tissue)^7^ routes, and is commonly characterized by multiple erythema migrans, appearing 3-5 weeks after the tick bite and usually smaller than the primary lesion^1^. Early disseminated Lyme disease is further commonly characterized by cranial nerve palsies, facial nerve palsies, and meningitis. Other symptoms typical of early disseminated Lyme disease include: fever, myalgia, arthralgia, headache, and fatigue. Finally, late Lyme disease occurs weeks to months after initial disease transmission, and the most common manifestation is arthritis, but may also involve encephalitis, encephalopathy, and polyneuropathy. Further, people affected by late Lyme disease may develop post-treatment Lyme disease syndrome (PTLDS), also known as chronic Lyme disease.

At present approximately 30,000 cases of Lyme disease are reported to the Center for Disease Control (CDC) each year by state health departments. However, the CDC estimates that the actual number of Americans diagnosed and treated for Lyme is closer to 300,000-476,000 people per year^8^. Further, Lyme is expanding, with more regions achieving high-incidence status for Lyme Disease^9^, as Lyme expands into new parts of the northern Midwest and New England regions. This spread is expected to worsen as more land is developed and chaparral is removed, and more climates become available for questing ticks owing to global climate-associated changes^9,10^.

As no currently-approved human vaccine exists, early detection and treatment remain the most effective methods to manage Lyme disease. Typical treatment for Lyme in the U.S. is a 100mg dose of Doxycycline taken twice daily for two weeks^11^. Generally objective manifestations of Lyme, such as erythema migrans, meningitis, or arthritis resolve during or after completion of a course of antibiotics, but some patients report long-term (*>*=6 months) persistence of fatigue, musculoskeletal pain, or difficulties with concentration and memory, which are thought to be PTLDS^11^. Typically Lyme is diagnosed by the presence of erythema migrans, clinical symptoms, and patient travel or activity in Lyme-endemic areas^11,12^. In the absence of erythema migrans, Lyme is diagnosed by a 2-tiered serologic test comprising an enzyme-linked immunoassay (ELISA) followed by a Western immunoblot. 2-tiered serologic tests, however, suffer from low sensitivity during the early disease stages, ranging anywhere from 26-89% depending on the specific test and test administrator, due largely to the time-lag in production of host antibodies^12^, which typically require a couple weeks to reach detectable levels. One study reports that the standard 2-tiered serology test missed 85.7% of cases of early Lyme disease with spirochetemia^13^.

Lack of a high-fidelity test for Lyme has led to misdiagnosis and heightened patient anxiety surrounding Lyme disease^1,12^. Further, misdiagnosis leads to over prescription of antibiotics, leading to potential increases in the public health burden of Lyme. Attempts to directly detect *Borrelia* spirochetes in patients have been proposed to increase clinical Lyme testing fidelity, primarily detection via blood culture^6,14^, polymerase chain reaction (PCR)^15,16^, a combination of the two^11,17,18^, or more recently aptamer technology^19^.

However, detection by culture methods has a relatively long turnaround time to results, requiring around 7 days to achieve greater than 90% detection rate^17^. Further, culture methods would be unable to detect non-cultivable infectious spirochetes, which have been found to persist following standard antibiotic treatment^20^ and make culture-based detection less efficacious. Additionally, PCR detection is unable to distinguish between live and dead *Borrelia*as it is simply amplifying *Borrelia*RNA or DNA^21^ making it an unsuitable for detecting active infection. Lastly, while promising, aptamer technology to this date detects *Borrelia*antigens rather than whole *Borrelia*^19^.

Direct detection via immunocapture however could overcome these shortfalls, as *Borrelia* could be captured on an immunofunctionalized surface directly from a peripheral blood draw. Further, immunocapture could detect *Borrelia* regardless of whether they are replicating, so long as they express an antigen corresponding to chosen capture antibodies. Of immunocapture devices, microfluidic chips are well-poised to capture *Borrelia* directly from peripheral blood, given their long history of capturing and detecting rare cells, such as capturing circulating tumor cells from whole blood^22,23^, detecting a variety pathogens such as viruses, bacteria, fungi, or parasites^24^, or rare-teratoma forming cells from differentiating stem cell populations^25^.

A major defining characteristic of Lyme disease is the drastic proteome changes *Borrelia* undergoes during infection^26^. In their natural environment, *Borrelia* are transmitted from one of the four species of hard tick within the I. Ricinus complex to a mammalian host^27^. The Lyme life cycle is thought to consist of spirochete-free tick larvae feeding on reservoir hosts: small mammals such as squirrels and mice, or birds^28,29^. Spirochete-free larvae acquire *Borrelia* infection by feeding on reservoir hosts and then pass *Borrelia* onto new reservoir hosts during the nymphal stage or larger mammals such as deer during the adult stage^27^, with humans thought to be a dead-end host. Discovery of this enzootic cycle led to a leading model in which outer surface protein A (OspA) is expressed by *Borrelia* in the tick midgut, then decreases in expression during tick bloodmeal, while another surface protein OspC increases in expression during bloodmeal, and then remains high during the early stages of host infection^26,30,31^. Understanding of this cycle has been critical to preventing the spread of Lyme, as early efforts led to the development of a vaccine against OspA, which has since been pulled from the market due to a lack of efficacy^26^. Similarly, we anticipate that engineering an immune-sensor for Lyme will require validating that sensor against *Borrelia* which are representative of *Borrelia* during the early stages of mammalian infection. In this paper we first report a culture system for recreating key Osp changes associated with Lyme, then describe a microfluidic immunocapture device for the direct detection of *Borellia burgdorferi*. We report that this chip could be used for microscopic detection via a fluorescent readout, or alternatively, as a platform to enrich spirochete density for PCR detection of *Borellia burgdorferi*.

## Methods

### Device fabrication and functionalization

Devices were fabricated by A.M. Fitzgerald & Associates (Burlingame, CA) according to previously described specifications^22,23^. The silicon device surfaces were functionalized with a previously described protocol using 3-mercaptopropyltrimethoxysilane (MPTMS) and N-gamma-maleimidobutyryloxysuccinimide ester (GMBS) NeutrAvidin-biotin chemistry (Thermo Fisher Scientific, Rockford, IL)^22,23^.The devices were functionalized with primary antibodies via a biotinylation linkage. NeutrAvidin-functionalized devices were incubated with 5 *μ*g/mL biotinylated rabbit anti-*Borrelia* anti-OspC antibody (Novus Biologicals, Littleton, CO). All antibodies and chip-functionalizations were prepared in 1% BSA in PBS. Polydimethysiloxane (PDMS) sheets (5:1 base:curing agent), approximately 3 mm thick, were polymerized for 18 hr at 60^*°*^C and trimmed to form covers for the GEDI chip. A PDMS sheet was clamped to the top of the post arrays. Inlet and outlet holes were created with a biopsy punch, and tygon tubing was inserted into the PDMS gasket to connect GEDI chip inlet to a syringe pump and outlet to a microcentrifuge tube for effluent collection. Devices were primed with a 50/50 Ethanol/DI water mixture, and then flushed with DI water water and PBS before experiments.

### 1.1 Cell Culture

The B31 *Borrelia* cell lines were obtained from ATCC (Manassas VA), product number: 35210. BxPC3 cells were obtained from ATCC (Manassas, VA). All cell lines were cultured in humidified incubators (37°C and 5% CO2, pH=7.0, or 27°C, and 1% CO2, pH=8.0) using BSK-H Medium with 6% rabbit serum (Millipore Sigma, Burlington, MA). pH was adjusted by adding 6M HCL (diluted from 37% HCL (Milipore Sigma, Burlington, MA)) or Sodium hydroxide pellets (Milipore Sigma, Burlington, MA). Media was supplemented with 100*μg/ml* DTT (VWR Life Sciences, Radnor, PA) and 50*μg/ml* Rifampicin (TCI, Montgomeryville PA). B31 were harvested after 3–7 days of culture, during the late-log to stationary growth phase.

### 1.2 Capture Experiments

Cells were centrifuged at 9800g for 10min directly from culture, and resuspended in PBS. Functionalized silicon devices were mounted with Tygon tubing inlets and outlets in a custom PMMA holder. Cell capture was achieved by flowing 0.66 mL of cell suspension through the device at 1 mL/h followed by effluent collection and a OD400 reading was obtained by distributing 200ml of either the effluent or input cell suspension into three wells of a 96-well plate, and reading optical density on a BioTek Synergy H1 plate reader.

### 1.3 Immunofluorescence

Cells were harvested from the late-log to stationary phase and centrifuged at 9,800g for 10 min in microcentrifuge tubes. All following steps were performed in microcentrifuge tubes. Culture media was aspirated and cells were then resuspended in PBS and centrifuged through the same cycle. Cells were then resuspended in 200-300*μL* of 4% PFA in PBS and incubated for 12 min. Excess PBS (>1ml) was used to dilute PFA solution and cells were centrifuged at 9,800g for 10min. The PFA-PBS supernatant was aspirated, and cells were resuspended in 1.5ml PBS. Cells were spun at 9,800g for 10 mins, and PBS supernatant was aspirated to remove residual 4% PFA. Cells were then either resuspended in 200-300*μL* 0.5% Saponin in PBS (for permeabilization stanis) or PBS, and incubated for 15mins at 37^*°*^C. Excess PBS was used to dilute permeabilization solution and cells were spun through the same cycle. A wash step with PBS followed by the same spin cycle was then performed. Cells were then blocked for 30 min at room temperature in 5% BSA in PBS. Blocking solution was removed by washing with excess PBS and spinning. Cells were then resuspended in an antibody solution containing 20*μL* 1% BSA in PBS, 5*μL* mouse anti-*Borrelia* OspA (LS Bio, Seattle, WA) (0.1mg/ml), 1*μL* of either rabbit anti-*Borrelia* OspC (Novus biologicals, Littleton, CO) (1mg/ml) or rabbit anti-*Borrelia* ErpN/OspE (Novus Biologicals, Littleton, CO) (1mg/ml). 2*μL* each of Secondary antibodies goat anti-rabbit IgG (H+L) Alexa Fluor Plus 555 (2mg/ml) and goat anti-mouse IgG (H+L) Alexa Fluor Plus 488 (2mg/ml) were added to this suspension. Antibodies were incubated overnight at 4^*°*^C for at least 18 hours. Control stains were performed similarly by replacing primary antibody volumes with a corresponding volume of 1% BSA. For experiments in which cells were stained on-chip, this antibody solution was pipeted onto dismounted GEDI chips after running, and allowed to incubate inside a humidifed chamber at 37^*°*^C for 1hr.

#### 1.3.1 CellProfiler Image Analysis

Quantitative data was obtained by first flat field correcting all images in Matlab. Phase contrast images were then processed in CellProfiler to create a mask. Total intensity of OspA and OspC or OspE was then measured inside the mask and reported as intensity per area occupied by *Borrelia*.

### 1.4 qRT-PCR

All qRT-PCR experiments were performed on a QuantStudio 3 Real-Time PCR system. A QuantiNova SYBR Green RT-PCR kit (Qiagen, Hilden Germany) was used to obtain cycle threshold (Ct) values. Samples were run in a total 20*μL* volume. Kit instructions form the manufacturer provided handbook were followed for all reagent volumes and cycling times, except for the denaturation, annealing and extension, and total cycles, which were 10s, 30s, and 45 cycles, respectively. Forward and reverse primers were purchased from Life Technologies Corporation (Carlsbad, CA). Forward primers for FlaB, OspA, OspC, and OspE are as follows: GATTAGCAGCGTTAATGC^32,33^, AAGTACGATCTAATTGCAACAGT^32,33^, CGGATTCTAATGCGGTTTTACTTG^7^, TGATGGGCAAAGTAATGGAGAGG^7^, respectively, Citations for primers are listed after the primer. Reverse primers can be obtained by reversing the text direction.

### 1.5 Statistics

All statistical tests were performed in GraphPad 8.0. For **figure 2** statistical tests were ordinary 1-way ANOVA across all genes and days of expression. For **figure 5** a Kruskal Wallis 1-way ANOVA was used for all days in OspA external experiments, M2 in OspC external experiments, and on all days in OspE external experiments. For all other tests (ie: M4/M7 in OspC external and all total expression experiments across all days for OspA, OspC, and OspE) in **figure 5** an ordinary 1-way ANOVA was used. Asterisks are as follows: *p<.05, **p<.01, ***p<.001, ****p<.0001. All data sets were tested for normality with a D’Agostino and Pearson test when n was large enough (OspA external and total IF experiments) or a Shapiro-Wilk test when n was not large enough (all other data sets).

## 2 Results

### Culture Validation

To recreate the Osp changes representative of early Lyme Disease we began by creating a culture system that mimics both tick and mammalian specific signals: temperature, pH, and atmospheric CO2 concentration (figure 1A). Previous studies have identified temperature^34–37^, pH of the culture medium^38^, atmospheric CO2 concentration^7^, or a combination of these factors^39–41^ as capable of recreating many of the proteomic changes seen in host-adapted spirochetes (spirochetes cultured in dialysis membrane chambers (DMCs) inside mice or rats). For our culture system we chose a pH of 8.0, a temperature of 27°C, and an atmospheric CO2 concentration of 1% as our tick-mimicking (TM) conditions and a pH of 7.0, a temperature of 37°C, and an atmospheric CO2 concentration of 5% as our mammalian-mimicking (MM) conditions. Upon culturing *Borrelia* in the TM and MM conditions we observed several *Borrelia* isoforms, such as round bodies (RBs), blebs, and colonies (figure 1B-F). All isoforms were observed in both TM and MM conditions at all time points, suggesting the environmental factors associated with TM or MM conditions do not induce their formation. These forms have been observed by other researchers^42,43^ and are thought to play a role in stress response, antibiotic resistance, and persistence within mammalian hosts.

**Figure 1.**
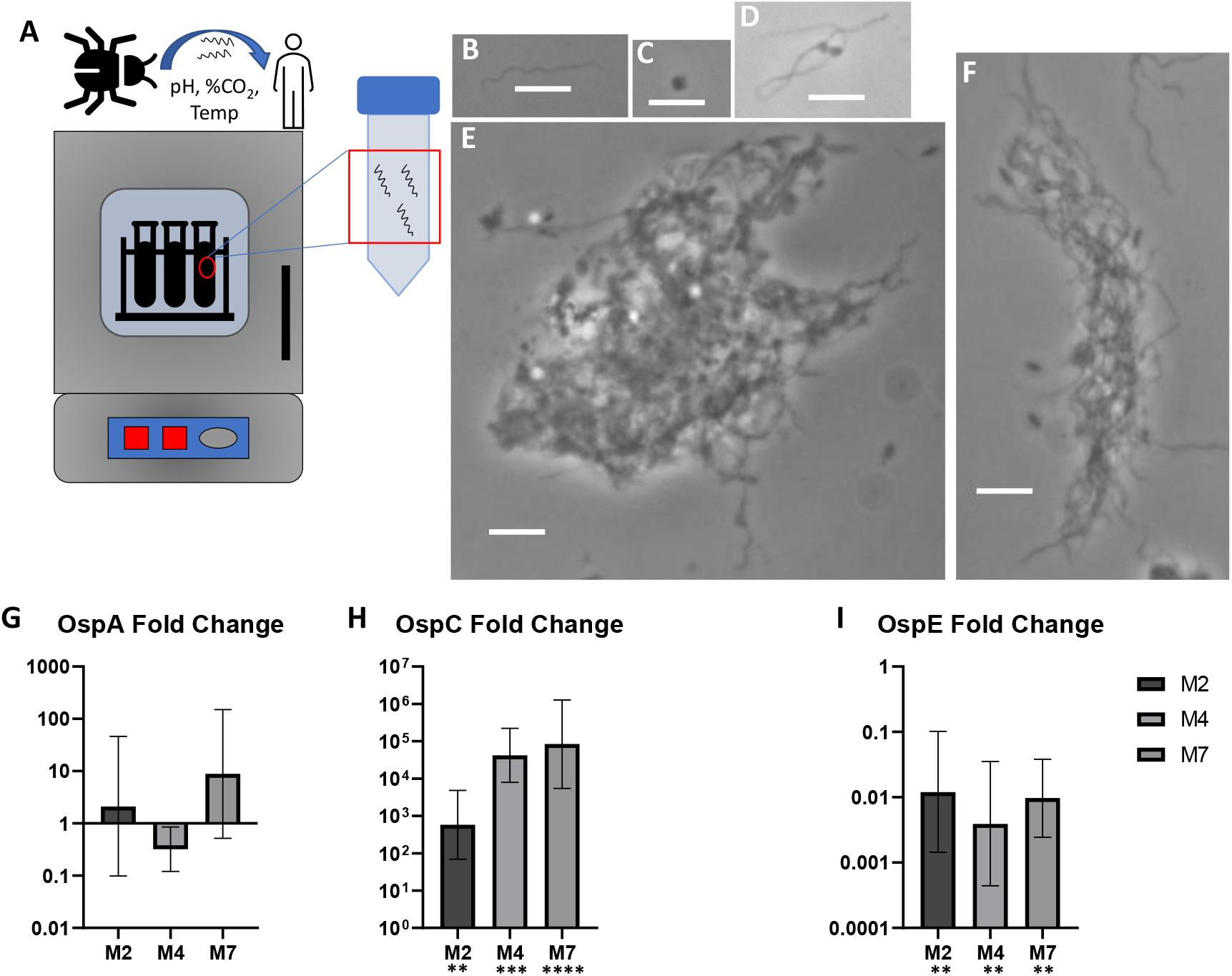
**A** Cartoon schematic depicting culture system designed to mimic environmental factors in the tick and mammalian environment. **B-F** Phase contrast images (100x) depicting *Borrelia* isoforms: spirochete (**B**), round body (**C**), bleb (**D**), and biofilm/colony (**E,F**). Scale bar is 5*μm*. **G-I** qRT-PCR results showing fold change gene expression relative to *Borrelia* cultured in TM conditions of OspA (n=4,3,5 respectively), OspC (n=3,3,6 respectively), and OspE (n=3, 3, 6 respectively), respectively, at 2, 4, and 7 days post transfer. (n=3 for all tick comparison data sets). Error bars show range of fold change gene expression for 1 standard deviation.

To confirm our culture system faithfully recreated Osp changes, we performed quantitative reverse-transcriptase polymerase chain reaction (qRT-PCR) experiments by culturing B31 *Borrelia* under TM conditions and then transferring *Borrelia* into MM conditions for either 2, 4, or 7 days. At the end of each time point, RNA was extracted from *Borrelia* whole-cell lysate using a Qiagen RNeasy Mini Kit and proteinase K. qRT-PCR was then performed on extracted RNA using a QuantiNova SYBR Green RT-PCR kit on a QuantStudio 3 Real-Time PCR system. Delta delta Ct analysis^44^ was then performed to determine fold-change gene expression data for OspA, OspC, and OspE at the 2, 4, and 7-day time points post transfer from the TM to MM conditions. Flagellin B (FlaB) was chosen as a control housekeeping gene as its expression has been shown to be relatively unaffected by temperature, pH, atmospheric CO2 concentration, or cultivation inside a DMC chamber^7,34,36,39,40^. Interestingly we notice a substantial upregulation of OspC and rather sporadic changes in OspA, with no consistent trends, and a down-regulation of OspE (figure 1G-I). Although the upregulation of OspC and no substantial changes in OspA were expected, the substantial down regulation of OspE is rather surprising as it is generally understood that OspE is not expressed by *Borrelia* in the tick nor during early infection, but is upregulated during the later stages of mammalian infection^26,31^. However, our results suggest that our culture system does recreate the dramatic upregulation of OspC seen during early Lyme infection, and indicates that OspC is likely a good antigen choice for immunocapture, particularly given the relatively few commercially available antibodies with known target antigens.

### Qualitative Immunofluorescence

Given that our qRT-PCR data indicates our culture system is recreating the population-level OspC upregulation which occurs during early Lyme disease, we sought to further investigate the behavior of individual spirochetes with phase contrast microscopy (PCM) and immunofluorescent (IF) imaging during transmission from TM to MM conditions. Spirochete heterogeneity is known to be a substantial aspect of *Borrelia* behavior. For example, *Ohnishi et al*^45^ found that *Borrelia* within unfed ticks were largely homogenous, with most producing only OspA. However, during bloodmeal many spirochetes begin producing OspC, leading to a heterogenous population with respect to OspA and OspC. Further, *Hefty et al*^46^ performed a set of experiments to show that not only does *Borrelia* regulate its surface protein expression in response environmental and host-specific factors, but also alters the cellular location of antigens during various phases of the enzootic cycle. To account for this expected heterogeneity in Osp expression we performed a set of experiments in which we stained *Borrelia* for either OspA and OspC, or OspA and OspE, then compared between samples in which *Borrelia* were surface stained (external) or permeabilized, and thus stained internally and externally (total expression). Interestingly, upon visual inspection we find *Borrelia* expressing either OspA, OspC, OspE, both OspA and OspC, both OspA and OspE, or neither (**figure 2** and **3**). We are able to find these subpopulations of *Borrelia* in either culture condition and at any time point, except MM conditions on day 7, and one group (OspA-/OspC-total stain) on day 4. Inspection of **figures 2** and **3** suggests an interesting trend: *Borrelia* become more homogenous in terms of OspA and OspC expression during later time points in mammalian conditions. Further, we note OspC expression, particularly in the externally stained samples, tends to be highly punctuated in expression. This could have implications for microfluidic device design, as the antigen-presenting surface must come into close contact with an antibody-functionalized surface.

**Figure 2.**
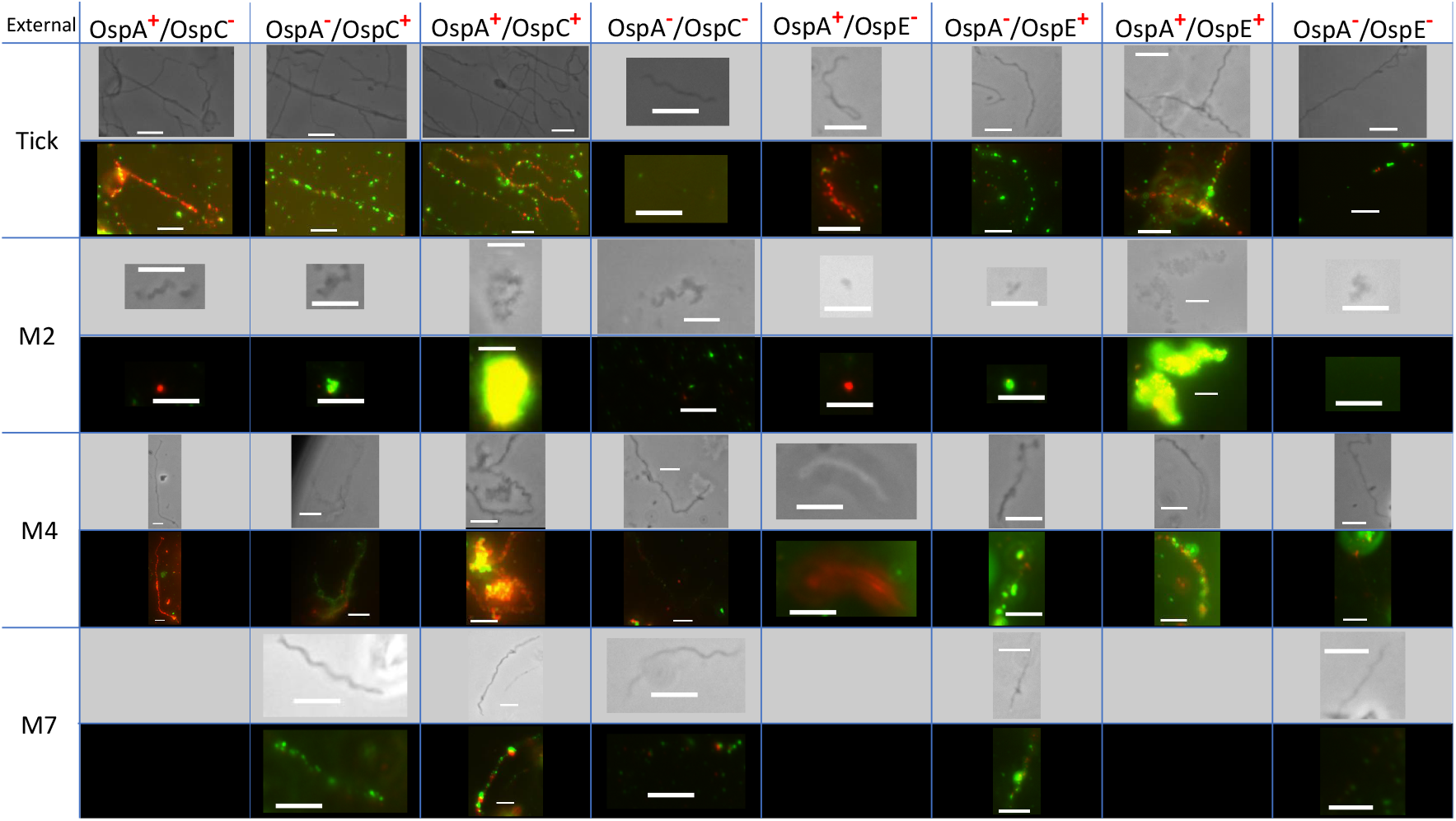
Phase contrast and corresponding immunofluorescent images (100x) showing expression of OspA and OspC, or OspA and OspE for externally stained *Borrelia*. Scale bar is 5*μm*. Note: OspC and OspE are shown in green, OspA is shown in red.

**Figure 3.**
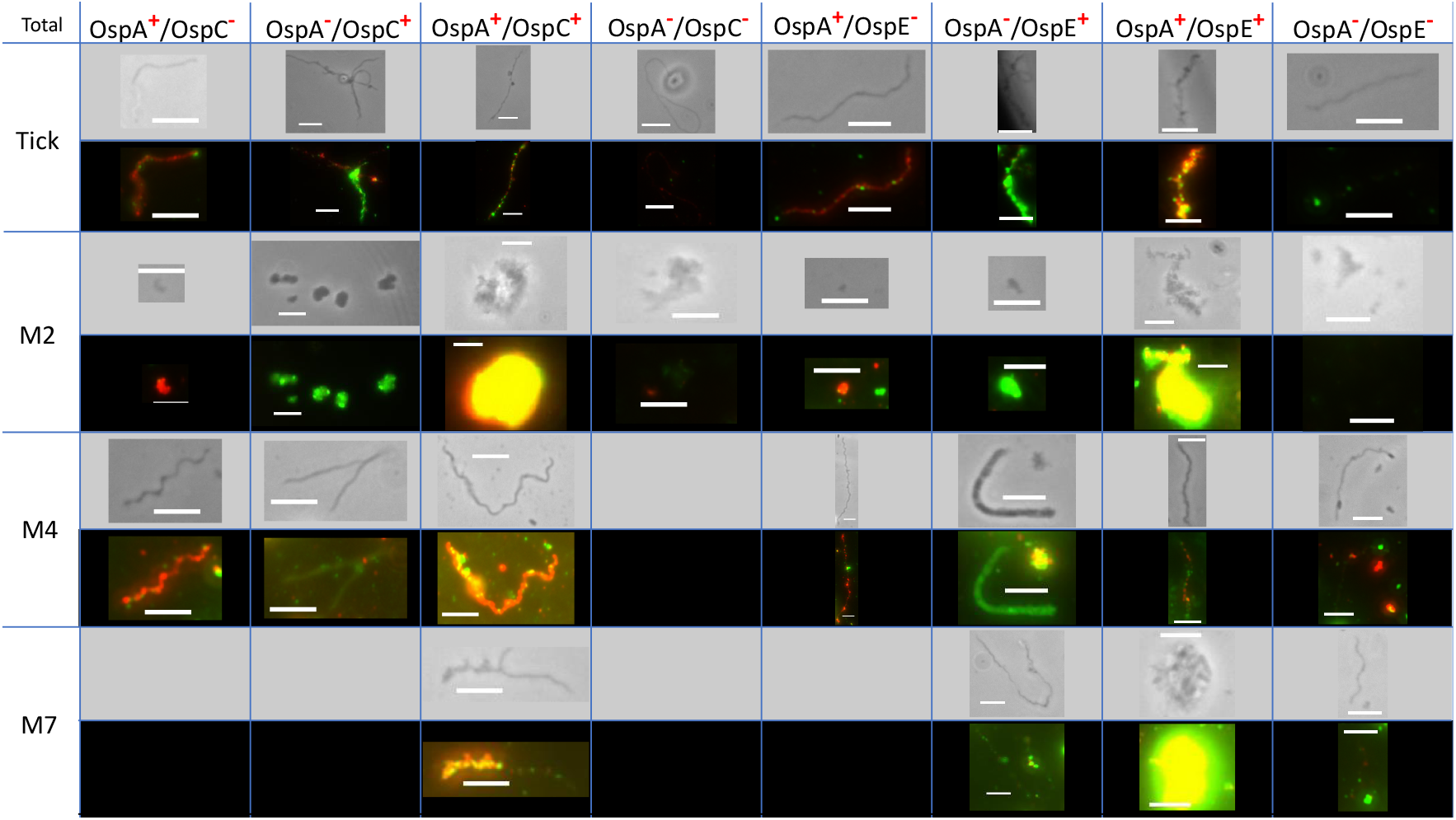
contrast and corresponding immunofluorescent images (100x) showing expression of OspA and OspC, or OspA and OspE for totally stained *Borrelia*Ṡcale bar is 5*μm*. Note: when possible a spirochete is shown instead of an isoform. Note: OspC and OspE are shown in green, OspA is shown in red.

Having investigated the heterogeneity of *Borrelia* as they transition from TM to MM conditions, we sought to investigate the heterogeneity of *Borrelia* pleomorphisms **(figure 4)**. In this investigation we sought primarily to determine which Osps were expressed externally, as only externally expressed antigens are relevant for capture. For our purposes we defined these pleomorphisms according to *Merilainen et al*^42^, in which blebs consist of a spirochete with a membrane bleb, RBs consist of a lone spherical structure, and Biofilms consist of a colony mostly of spirochetes, but containing blebs and RBs.

**Figure 4.**
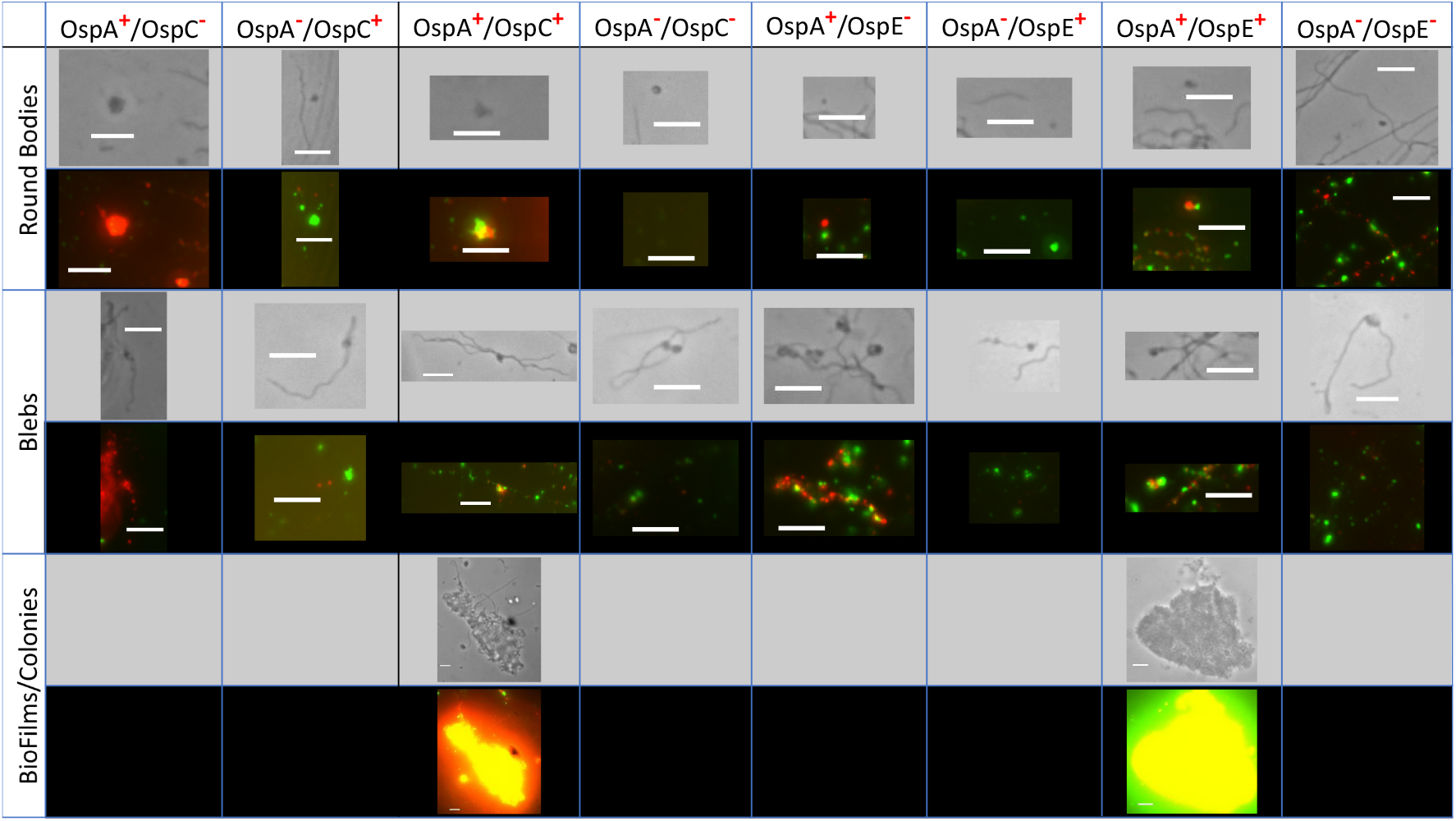
Phase contrast and corresponding immunofluorescence images (100x) showing expression of OspA and OspC, or OspA and OspE for externally stained *Borrelia* isoforms. Scale bar is 5*μ* m. Note: OspC and OspE are shown in green, OspA is shown in red.

Strikingly, we observe that the Bleb and RB forms express all variations of *OspA*^+*/*−^*/OspC*^+*/*−^ and *OspA*^+*/*−^*/OspE*^+*/*−^, while BioFilms/Colonies were only found to express both markers, likely due to the large number of *Borrelia* making up the colony.

### Quantitative Immunofluorescence

Having qualitatively analyzed our IF images, we sought to perform a quantitative investigation of how Osp expression changes during the first week of transition from TM to MM conditions. As individual spirochetes often overlap or are entangled with several others, and given individual spirochetes are often impossible to single out within colonies, we calculated an intensity per *Borrelia* area for each image. An overview of our image processing pipeline is shown in figure 5a. In brief, it consists of taking the phase contrast (PC) image, performing a flat field correction, followed by a number of image processing algorithms which result in a mask of the PC image. Each of the OspA, OspC, or OspE IF images is then multiplied by the mask so that fluorescent signal is only counted when a *Borrelia* (in any isoform) is present. Thus our image analysis shows the intensity per *Borrelia* area for each image, and does not bias against *Borrelia* with low or no expression of Osps, as masks are created from PC images. Strikingly, our results are in agreement with qRT-PCR data for OspA, as there is no consistent trend in OspA expression change during the 7 day time course. For OspE our results show little change in expression between days in MM conditions, but do not show a down-regulation relative to TM conditions. Interestingly, for OspC, we see a noticeable upregulation in OspC expression during transition from TM to MM conditions, but only for totally stained *Borrelia* and not externally stained *Borrelia* which suggests immunocapture with OspC may not necessarily be made easier by upregulation of OspC expression. However, given our qRT-PCR data suggests a strong upregulation of OspC, and OspC expression is found under all conditions at all time points, and in all *Borrelia* isoforms, we chose OspC as our target capture antibody.

**Figure 5.**
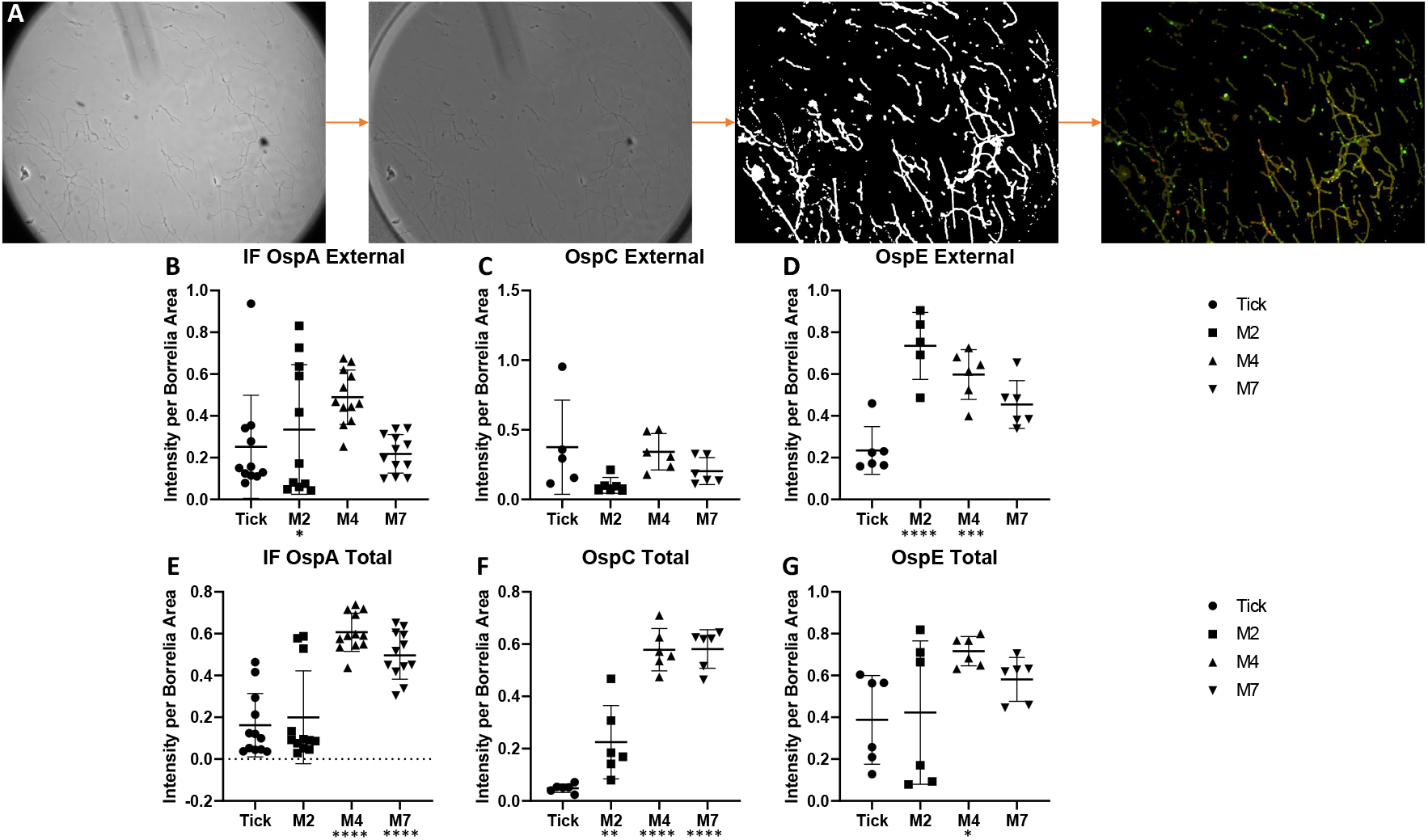
**A** Schematic depicting image analysis pipeline: raw images were flat field corrected, then masked to identify locations where *Borrelia* are present. Masked images are then multiplied by IF images so that signal is only counted in locations where *Borrelia* are present. **B-D** Graphs depicting intensity per total *Borrelia* area for OspA, OspC, and OspE. Note: each dot represents one single image (taken at 100x). Time post transfer refers to time in MM conditions after being transferred from TM conditions.

### Immunocapture

Having decided on OspC as our target capture antibody, we first began by testing capture on a static surface in which minimal shear force is present. To perform this test *Borrelia* were seeded onto an anti-OspC functionalized glass coverslips, and allowed to bind for 5 minutes. Slides were gently washed in PBS and then imaged with PC microscopy for detection of *Borrelia*. As seen in **figure 6d** numerous *Borrelia* are found on the anti-OspC surface. Comparison to a surface functionalized with an isotype control antibody resulted in a blank coverslip (data not shown), demonstrating specificity of capture.

**Figure 6.**
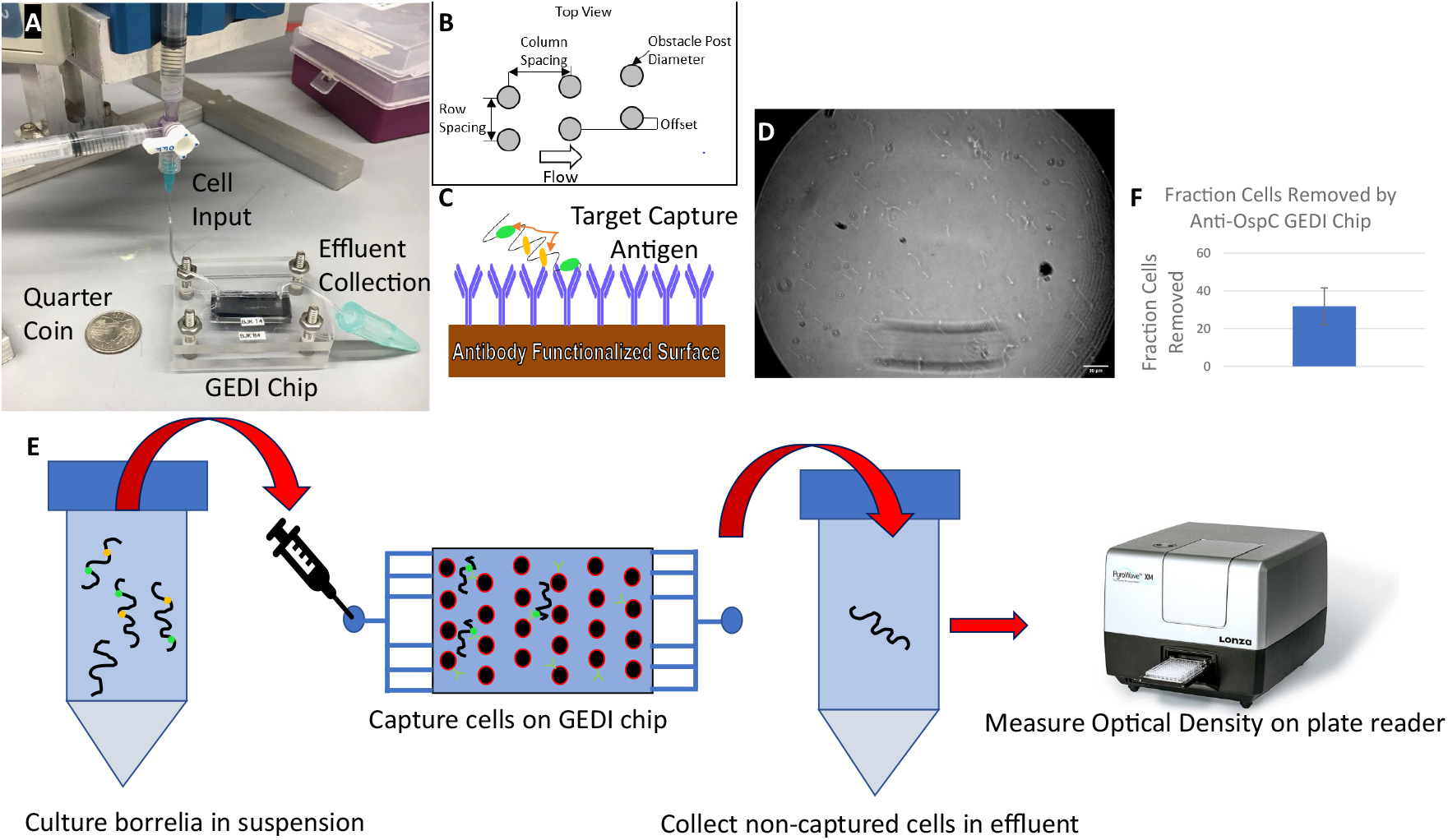
**A** Picture of GEDI chip on lab bench, note quarter for scale. **B** Schematic depicting GEDI obstacle array geometry. **C** Schematic depicting *Borrelia* capture with immuno-functionalized surface. **D** Phase contrast image of anti-OspC functionalized glass coverslip demonstrating *Borrelia* capture (40x). Scale bar is 30*μm*. **E** Cartoon schematic depicting process for determining removal efficiency. *Borrelia* are cultured in suspension, pumped through GEDI device, effluent is collected, and OD400 is read on plate reader. **F** Graph showing fraction of cells removed by GEDI device.

Having confirmed that the anti-OspC antibody is not limited by steric hindrance or lack of a readily available epitope, we began testing our microfluidic device’s performance. The microfluidic immunocapture device is termed geometrically enhanced differential immunocapture (GEDI). In short, the device is a Hele-Shaw cell with a staggered obstacle array. The principle idea driving device design is that the array geometry is designed to maximize the number of collisions particles (in this case *Borrelia*) experience with an antibody-functionalized post (**figure 6b**), thus ensuring all *Borrelia* end up on a streamline which brings them in close contact with an antibody-functionalized surface. Further, the posts serve to maximize the available capture area, by drastically increasing the surface-to-volume ratio inside the microfluidic chip. For a more detailed description of device design see:^47,48^.

To evaluate device performance, we began by taking a suspension of *Borrelia* culture and pumping that suspension through our device with an infusion pump. The effluent was then collected in a microcentrifuge tube and the optical density at 400nm (OD400) was measured on a Lonza PlateReader (**figure 6a** and **e**). By comparing the OD400 of the input cell suspension and effluent from the device, we calculated the fraction of cells removed by the GEDI chip. Our results show that on average the anti-OspC GEDI device removes 31.8% of *Borrelia* from a culture suspension grown to late-log phase under MM conditions. After evaluating capture performance we sought to investigate Osp expression of *Borrelia* captured on chip (**figure 7b**).

**Figure 7.**
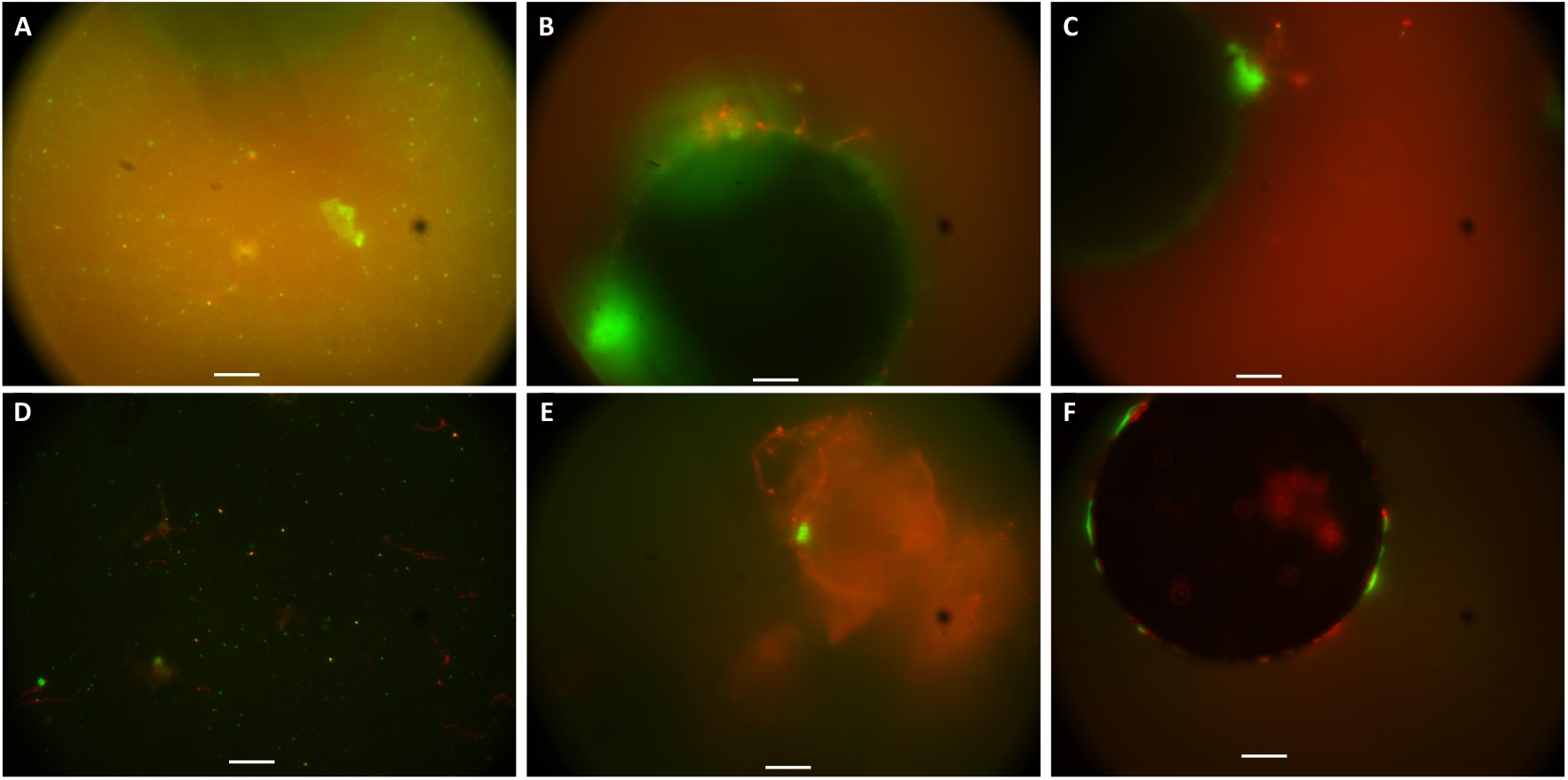
On-chip IF images showing **A** *Borrelia* RBs expressing OspC and or OspA, **B-C** Capture of OspC and OspA expressing spirochetes on obstacle post, **D** capture of *Borrelia* RBs expressing OspE, **E** capture of *Borrelia* biofilms expressing OspA and OspE, and **F** Capture of spirochetes expressing OspA or OspE. Note: images are 100x, scale bar is 5*μm*. Top images are OspC (green) and OspA (red), bottom images are OspE (green) and OspA (red).

Interestingly we see spirochetes bound to antibody functionalized posts (**figure 7b-c**) expressing OspC near the post, with OspA expression along the body of the spirochete (where fluid forces would pull the OspC-free spirochete body away from the post). Further, we notice our device captures a substantial number of RB *Borrelia* along the bottom surface, suggesting RBs settle to the bottom of the device due to gravitational forces, and then are captured by anti-OspC antibodies as they roll along the bottom surface (figure 7a). Further, we found our device is also capable of capturing biofilms (figure 7e) expressing a combination of OspA and OspE. We also identify both RBs and spirochetes expressing solely OspA, OspE, or both when we stained collectively for OspA and OspE. Thus on-chip imaging demonstrates our device is capable of capturing *Borrelia* isoforms expressing one of the OspA, OspC, or OspE proteins, or combination of OspA and OspC or OspA and OspE.

## 3 Discussion

In designing a culture system that would accurately recreate Osp expression associated with the tick-to-mammalian transmission of *Borrelia* during early Lyme Disease, we chose the three environmental factors most commonly discussed in the literature which recreate the Osp changes that occur during the tick-to-mammal transition: temperature^34–37^, pH of culture media^38^, and atmospheric CO2 concentration^7^. Although the literature on this topic varies with regards to exact test conditions, there is a general trend that conditions closer to those of mammalian blood (ie: 35-37C, pH =7, and 5% dissolved CO2) cause an upregulation of OspC and OspE, and either little change or down-regulation of OspA. It is interesting that our qRT-PCR results are in agreement with these trends for OspA and OspC, but are in disagreement with existing data on OspE. One possible explanation for this is that previous studies^36^ have found that while a temperature shift did induce an upregulation of OspE, OspE was down-regulated in fed ticks, suggesting host-specific signals can override the positive stimulus provided by temperature^36^, which could be the case for the combined effect of temperature, pH, and CO2 concentration in our culture system. In summary however, our culture system does recreate the upregulation of OspC seen from antibody response studies during early infection of canines and mice with *Borrelia*^31,36^.

Investigation of our IF images reveals substantial heterogeneity in expression of Osp proteins on individual *Borrelia*. This is in line with previous studies which have found similar heterogeneity in Osp expression among *Borrelia* whether they are cultured in vitro, in DMCs, or collected from ticks^45,46^. It is interesting to note previous studies have found the same beaded expression of OspE^**?**^ as we did but did not find the same beaded expression we observed for OspC. The cause of this heterogeneity *in vivo* is thought to arise from the role these proteins play in movement of *Borrelia* from the unfed tick midgut to the tick salivary glands during feeding. *Ohnishi et al*^45^ propose a model in which non-infectious *Borrelia* expressing neither OspA or OspC, or solely OspA infect the host dermis early on during tick feeding, with *Borrelia* expressing neither OspA or OspC, or a subgroup expressing only OspC establishing stable infection in the host dermis later on. However, *Grimm et al*^49^ performed a series of studies with a mutant *Borrelia* strain lacking OspC and showed OspC was required for stable infection of mice, regardless of infection route (needle or tick). A possible explanation for this discrepancy is that *Ohnishi et al*^45^ investigated the mouse dermis attached to the tick hypostome rather than attempting to retrieve infectious *Borrelia* from the mouse through feeding ticks. The OspA and OspC double negative *Borrelia* detected in mouse dermis could then represent a non-infectious population of spirochetes, which would be in line with the presence of these double negative spirochetes in mice that were not stably infected^45^.

These findings have a substantial impact on the design of immunocapture devices as we chose OspC as our primary capture antigen due to available data demonstrating OspC sero-reactivity across multiple mammalian species^31,45^, and although there is sero-reactivity in humans to OspC^50^ behavior of *Borrelia* in humans often differs from model organisms such as mice, due to humans not being part of the natural enzootic cycle of *Borrelia*. However, if future studies demonstrate a substantial fraction of infectious *Borrelia* expressing Osps other than OspC during early infection, our device performance could be enhanced by functionalizing the device with a combination of multiple *Borrelia*-specific antibodies. Further, the beaded expression pattern we observed for OspC has implications for chip design, as it suggests that not only do *Borrelia* need to be brought in close contact with an antibody-functionalized surface, but that device performance might be enhanced by mixing of the cell suspension, so that the chance the OspC expressing surface of the *Borrelia* is brought into contact with the antibody-functionalized surface is maximized. One method of achieving this may be through incorporating the staggered herringbone mixer design developed by *Stroock et al*^51^ in our GEDI device.

Of further interest is our results showing expression of OspA, OspC, and OspE among some RBs and blebs, as well as biofilms. Although we are unaware of any literature on the expression of these Osp proteins by *Borrelia* isoforms, our data is encouraging from an immunocapture perspective as on-chip imaging demonstrates that our device captures *Borrelia* in any isoform. These isoforms are thought to largely arise from a *Borrelia* stress response, particularly due to antibiotics^42,43^. Our ability to capture these isoforms is promising as it is plausible *Borrelia* exist in one or multiple of these forms during infection in humans as they are entering into a stressful foreign environment, and would be exposed to antibiotics if the patient is actively being treated for Lyme^43^.

## 4 Conclusion

In summary, our device is the first reported immunocapture device for the detection of whole-organism *Borrelia*. Although our *Borrelia* removal rate was only 31.8%, this removal rate is expected to be sufficient for positive identification of *Borrelia*. Previous studies have shown that only 78%^36^ of *Borrelia* spirochetes cultured under similar conditions express OspC in vitro. Further, unbeknownst to us at the time of experimentation, OspC expression has been shown to decrease upon repeated passage of *Borrelia* at 37^*°*^C^30^. As our capture experiments were performed on *Borrelia* passaged through MM conditions for 2-4 passages, we deem our experiments an appropriate challenge to the system’s ability to capture *Borrelia* under a variety of expression conditions.

Further, although this work’s results focused on IF assays to detect *Borrelia* captured on chip, our device could serve as a platform to enhance the concentration of *Borrelia* present in patient blood into a small volume, which could be combined with work by other researchers to detect *Borrelia* with PCR^16^. This approach to Lyme detection would be particularly useful given the presence of non-cultivable spriochetes in antibiotic treated mice^20,52^. While our work lays the ground work for microfluidic immunocapture and detection of Lyme, future work should focus on three things: a multi-antibody functionalization scheme to address the heterogeneity during Lyme infection, microfluidic mixing to ensure non-uniformly expressed antigens come into contact with antibody-functionalized surfaces, and development of an on-chip PCR assay to detect captured *Borrelia*. Lastly, our microfluidic chip could be used in conjunction with aptamer technology as an alternative to immunocapture to detect either *Borrelia*antigen or potentially whole-cell *Borrelia*.

## 5 Data Availability

The data that support this study are available from the corresponding author upon reasonable request

## Author contributions statement

All experiments were performed by K.W. Writing was done by K.W. and B.K. All work was carried out at Cornell University, the second, current institute for K.W. is added for easier contact.

## References

1. Murray, T. S. & Shapiro, E. D. Lyme disease. Clin. Lab. Medicine 30, 311–328, DOI: 10.1016/j.cll.2010.01.003 (2010).

2. Piesman, J., Mather, T. N., Sinsky, R. J. & Spielman, A. Duration of tick attachment and Borrelia burgdorferi transmission. J. Clin. Microbiol. 25, 557–558, DOI: 10.1128/jcm.25.3.557-558.1987 (1987).

3. England, T. N. of Lyme Disease After an Ixodes Scapularis Tick Bite. Engl. J. 345, 79–84 (2001).

4. Gerber, M., Shapiro, E., Burke, G., Parcells, V. & Bell, G. The New England Journal of Medicine LYME DISEASE IN CHILDREN IN SOUTHEASTERN CONNECTICUT. (1996).

5. England, T. N. Journal Medicine ©. 209–215 (1998).

6. Wormser, G. P. Hematogenous dissemination in early Lyme disease. Wiener Klinische Wochenschrift 118, 634–637, DOI: 10.1007/s00508-006-0688-9 (2006).

7. Hyde, J. A., Trzeciakowski, J. P. & Skare, J. T. Borrelia burgdorferi alters its gene expression and antigenic profile in response to CO2 levels. J. Bacteriol. 189, 437–445, DOI: 10.1128/JB.01109-06 (2007).

8. CDC Lyme Data and Surveillance kernel description. https://www.cdc.gov/lyme/datasurveillance/index.html. xAccessed: 2021-07-16.

9. Kugeler, K. J., Farley, G. M., Forrester, J. D. & Mead, P. S. Geographic distribution and expansion of human lyme disease, United States. Emerg. Infect. Dis. 21, 1455–1457, DOI: 10.3201/eid2108.141878 (2015).

10. Stone, B. L., Tourand, Y. & Brissette, C. A. Brave New Worlds: The Expanding Universe of Lyme Disease. Vector-Borne Zoonotic Dis. 17, 619–629, DOI: 10.1089/vbz.2017.2127 (2017).

11. Stanek, G., Wormser, G. P., Gray, J. & Strle, F. Lyme borreliosis. The Lancet 379, 461–473, DOI: 10.1016/S0140-6736(11)60103-7 (2012).

12. Moore, A., Nelson, C., Molins, C., Mead, P. & Schriefer, M. Current guidelines, common clinical pitfalls, and future directions for laboratory diagnosis of lyme disease, United States. Emerg. Infect. Dis. 22, 1169–1177, DOI: 10.3201/eid2207.151694 (2016).

13. Lee, S. H. et al. Dynamic methylation and expression of Oct4 in early neural stem cells. J. Anat. 217, 203–213, DOI: 10.1111/j.1469-7580.2010.01269.x (2010).

14. Sapi, E. et al. Improved culture conditions for the growth and detection of Borrelia from human serum. Int. J. Med. Sci. 10, 362–376, DOI: 10.7150/ijms.5698 (2013).

15. Karan, L. et al. Dynamics of spirochetemia and early PCR detection of Borrelia miyamotoi. Emerg. Infect. Dis. 24, 860–867, DOI: 10.3201/eid2405.170829 (2018).

16. Lee, S. H., Vigliotti, V. S., Vigliotti, J. S., Jones, W. & Pappu, S. Increased sensitivity and specificity of Borrelia burgdorferi 16S ribosomal DNA detection. Am. J. Clin. Pathol. 133, 569–576, DOI: 10.1309/AJCPI72YAXRHYHEE (2010).

17. Liveris, D. et al. Improving the yield of blood cultures from patients with early lyme disease. J. Clin. Microbiol. 49, 2166–2168, DOI: 10.1128/JCM.00350-11 (2011).

18. Coulter, P. et al. Two-year evaluation of Borrelia burgdorferi culture and supplemental tests for definitive diagnosis of lyme disease. J. Clin. Microbiol. 43, 5080–5084, DOI: 10.1128/JCM.43.10.5080-5084.2005 (2005).

19. Kamarudin, N. A. A. N., Mawang, C. I. & Ahamad, M. Direct detection of lyme borrelia: Recent advancement and use of aptamer technology, DOI: 10.3390/biomedicines11102818 (2023).

20. Barthold, S. W. et al. Ineffectiveness of tigecycline against persistent Borrelia burgdorferi. Antimicrob. Agents Chemother. 54, 643–651, DOI: 10.1128/AAC.00788-09 (2010).

21. Li, X. et al. Burden and viability of borrelia burgdorferi in skin and joints of patients with erythema migrans or lyme arthritis. Arthritis Rheum. 63, 2238–2247, DOI: 10.1002/art.30384 (2011).

22. Gleghorn, J. P. et al. Capture of circulating tumor cells from whole blood of prostate cancer patients using geometrically enhanced differential immunocapture (GEDI) and a prostate-specific antibody. Lab on a Chip 10, 27–29, DOI: 10.1039/b917959c (2010).

23. Kirby, B. J. et al. Functional characterization of circulating tumor cells with a prostate-cancer-specific microfluidic device. PLoS ONE 7, 1–10, DOI: 10.1371/journal.pone.0035976 (2012).

24. Steven M. Singer#, Marc Y. Fink, V. V. A. HHS Public Access. Physiol. & behavior 176, 139–148, DOI: 10.1016/j.bios.2018.05.050.Point-of-care (2019).

25. Wellmerling, K., Lehmann, C., Singh, A. & Kirby, B. J. Microfluidic chip for label-free removal of teratoma-forming cells from therapeutic human stem cells. J. Immunol. Regen. Medicine 10, 100030, DOI: 10.1016/j.regen.2020.100030 (2020).

26. Manuscript, A. & Proteins, O. S. P. O Box 1842 Bushbuckridge 27 March 2018 APPLICATION FOR INTERN – ENVIRONMENT AND CLIMATE CHANGE PROGRAMME :. 2018, DOI: 10.1111/j.1574-695X.2012.00980.x.The (2018).

27. Radolf, J. D., Caimano, M. J., Stevenson, B. & Hu, L. T. Of ticks, mice and men: Understanding the dual-host lifestyle of Lyme disease spirochaetes. Nat. Rev. Microbiol. 10, 87–99, DOI: 10.1038/nrmicro2714 (2012).

28. Levine, J. F., Wilson, M. L. & Spielman, A. Mice as reservoirs of the Lyme disease spirochete. Am. J. Trop. Medicine Hyg. 34, 355–360, DOI: 10.4269/ajtmh.1985.34.355 (1985).

29. Bosler, E. M., Ormiston, B. G., Coleman, J. L., Hanrahan, J. P. & Benach, J. L. Prevalence of the Lyme disease spirochete in populations of white-tailed deer and white-footed mice. Yale J. Biol. Medicine 57, 651–659 (1984).

30. Schwan, T. G. & Piesman, J. Temporal changes in outer surface proteins A and C of the lyme disease-associated spirochete, Borrelia burgdorferi, during the chain of infection in ticks and mice. J. Clin. Microbiol. 38, 382–388, DOI: 10.1128/jcm.38.1.382-388.2000 (2000).

31. Wagner, B. et al. Antibodies to Borrelia burgdorferi OspA, OspC, OspF, and C6 antigens as markers for early and late infection in dogs. Clin. Vaccine Immunol. 19, 527–535, DOI: 10.1128/CVI.05653-11 (2012).

32. Persing, D. H., Telford, S. R., Spielman, A. & Barthold, S. W. Detection of Borrelia burgdorferi infection in Ixodes dammini ticks with the polymerase chain reaction. J. Clin. Microbiol. 28, 566–572, DOI: 10.1128/jcm.28.3.566-572.1990 (1990).

33. Kawabata, H., Tashibu, H., Yamada, K., Masuzawa, T. & Yanagihara, Y. Polymerase Chain Reaction Analysis of Borrelia Species Isolated in Japan. Microbiol. Immunol. 38, 591–598, DOI: 10.1111/j.1348-0421.1994.tb01828.x (1994).

34. Brooks, C. S., Vuppala, S. R., Jett, A. M. & Akins, D. R. Identification of Borrelia burgdorferi outer surface proteins. Infect. Immun. 74, 296–304, DOI: 10.1128/IAI.74.1.296-304.2006 (2006).

35. Brooks, C. S., Hefty, P. S., Jolliff, S. E. & Akins, D. R. Global analysis of Borrelia burgdorferi genes regulated by mammalian host-specific signals. Infect. Immun. 71, 3371–3383, DOI: 10.1128/IAI.71.6.3371-3383.2003 (2003).

36. Hefty, P. S. et al. Regulation of OspE-related, OspF-related, and Elp lipoproteins of Borrelia burgdorferi strain 297 by mammalian host-specific signals. Infect. Immun. 69, 3618–3627, DOI: 10.1128/IAI.69.6.3618-3627.2001 (2001).

37. Obonyo, M., Munderloh, U. G., Fingerle, V., Wilske, B. & Kurtti, T. J. Borrelia burgdorferi in tick cell culture modulates expression of outer surface proteins A and C in response to temperature. J. Clin. Microbiol. 37, 2137–2141, DOI: 10.1128/jcm.37.7.2137-2141.1999 (1999).

38. Carroll, J. A., Garon, C. F. & Schwan, T. G. Effects of environmental pH on membrane proteins in Borrelia burgdorferi. Infect. Immun. 67, 3181–3187, DOI: 10.1128/iai.67.7.3181-3187.1999 (1999).

39. Revel, A. T., Talaat, A. M. & Norgard, M. V. DNA microarray analysis of differential gene expression in Borrelia burgdorferi, the Lyme disease spirochete. Proc. Natl. Acad. Sci. United States Am. 99, 1562–1567, DOI: 10.1073/pnas.032667699 (2002).

40. Ramamoorthy, R. & Scholl-Meeker, D. Borrelia burgdorferi proteins whose expression is similarly affected by culture temperature and pH. Infect. Immun. 69, 2739–2742, DOI: 10.1128/IAI.69.4.2739-2742.2001 (2001).

41. Yang, X. et al. Interdependence of environmental factors influencing reciprocal patterns of gene expression in virulent Borrelia burgdorferi. Mol. Microbiol. 37, 1470–1479, DOI: 10.1046/j.1365-2958.2000.02104.x (2000).

42. Meriläinen, L., Herranen, A., Schwarzbach, A. & Gilbert, L. Morphological and biochemical features of Borrelia burgdorferi pleomorphic forms. Microbiol. (United Kingdom) 161, 516–527, DOI: 10.1099/mic.0.000027 (2015).

43. Rudenko, N., Golovchenko, M., Kybicova, K. & Vancova, M. Metamorphoses of Lyme disease spirochetes: Phenomenon of Borrelia persisters. Parasites Vectors 12, 1–10, DOI: 10.1186/s13071-019-3495-7 (2019).

44. Livak, K. J. & Schmittgen, T. D. Analysis of relative gene expression data using real-time quantitative PCR and the 2-ΔΔCT method. Methods 25, 402–408, DOI: 10.1006/meth.2001.1262 (2001).

45. Ohnishi, J., Piesman, J. & De Silva, A. M. Antigenic and genetic heterogeneity of Borrelia burgdorferi populations transmitted by ticks. Proc. Natl. Acad. Sci. United States Am. 98, 670–675, DOI: 10.1073/pnas.98.2.670 (2001).

46. Hefty, P. S., Jolliff, S. E., Caimano, M. J., Wikel, S. K. & Akins, D. R. Changes in temporal and spatial patterns of outer surface lipoprotein expression generate population heterogeneity and antigenic diversity in the lyme disease spirochete, Borrelia burgdorferi. Infect. Immun. 70, 3468–3478, DOI: 10.1128/IAI.70.7.3468-3478.2002 (2002).

47. Smith, J. P., Lannin, T. B., Syed, Y. A., Santana, S. M. & Kirby, B. J. Parametric control of collision rates and capture rates in geometrically enhanced differential immunocapture (GEDI) microfluidic devices for rare cell capture. Biomed. Microdevices 16, 143–151, DOI: 10.1007/s10544-013-9814-4 (2014).

48. Gleghorn, J. P., Smith, J. P. & Kirby, B. J. Transport and collision dynamics in periodic asymmetric obstacle arrays: Rational design of microfluidic rare-cell immunocapture devices. Phys. Rev. E - Stat. Nonlinear, Soft Matter Phys. 88, 1–9, DOI: 10.1103/PhysRevE.88.032136 (2013).

49. Grimm, D. et al. Outer-surface protein C of the Lyme disease spirochete: A protein induced in ticks for infection of mammals. Proc. Natl. Acad. Sci. United States Am. 101, 3142–3147, DOI: 10.1073/pnas.0306845101 (2004).

50. Anguita, J., Hedrick, M. N. & Fikrig, E. Adaptation of Borrelia burgdorferi in the tick and the mammalian host. FEMS Microbiol. Rev. 27, 493–504, DOI: 10.1016/S0168-6445(03)00036-6 (2003).

51. Stroock, A. D. et al. Chaotic Mixer for Microchannels Published by : American Association for the Advancement of Science Stable URL: http://www.jstor.org/stable/3075690 REFERENCES Linked references are available on JSTOR for this article : You may need to log in to JSTOR to a. 295, 647–651 (2016).

52. Hodzic, E., Imai, D., Feng, S. & Barthold, S. W. Resurgence of persisting non-cultivable Borrelia burgdorferi following antibiotic treatment in mice. PLoS ONE 9, DOI: 10.1371/journal.pone.0086907 (2014).

